# Mechanics of inactive swelling and bursting of porate pollen grains

**DOI:** 10.1101/2021.05.24.445387

**Authors:** Anže Božič, Antonio Šiber

**Affiliations:** Department of Theoretical Physics, Jožef Stefan Institute, 1000 Ljubljana, Slovenia; Institute of Physics, 10000 Zagreb, Croatia

## Abstract

The mechanical structure of pollen grains, typically characterized by soft apertures in an otherwise stiff exine shell, guides their response to changes in the humidity of the environment. These changes can lead both to infolding but also to excessive swelling and even bursting of pollen grains. We use an elastic model to explore the mechanics of pollen grain swelling and the role that soft, circular apertures (pores) play in this process. We identify and explore a mechanical weakness of the pores, which are prone to a rapid inflation once the grain swells to a critical extent. This transition leads to the bursting of the grain and the release of its content. Our results shed light on the inactive part of the mechanical response of pollen grains to hydration once they land on a stigma as well as on bursting of airborne pollen grains during rapid changes in air humidity.

## I. INTRODUCTION

The ability of pollen grains to adapt to physical and (bio-)chemical changes in their environment is essential for their survival. Once pollen grains leave the environment of the anther, they start to lose water, and this process activates a variety of protective mechanisms [1, 2]. Upon landing on a stigma, they take in exudate from its cells, swell, and eventually germinate, forming a pollen tube through which they fertilize the flower [3, 4]. While the growth of the pollen tube is an active response of the pollen grain, which requires coherent mobilization of numerous cellular mechanisms, a precise regulation of osmotic forces [5], and deposition of bio-polymers in the growing tube [6, 7], the response of the pollen grains to changes in their water content is, to a large degree, inactive and a consequence of their mechanical constitution [8–10]. This mechanical makeup has its drawbacks, however, as it can induce an extreme response of the grains in non-reproductive context—for instance, when the grains have not yet reached the stigma and the delicate chemical balance required for pollen tube growth has not been established. When the conditions in the environment change suddenly, e.g., when the relative humidity of the atmosphere increases significantly, such a response may lead to the bursting of pollen grains [9, 11]. This typically occurs in wind-borne pollen grains once they have been lifted up from the anthers into the atmosphere. For instance, it has been demonstrated that the concentration of grass allergens in non-pollen-containing fraction of ambient air correlates with air humidity [12], which has in turn been related to the bursting of pollen grain [11].

Understanding pollen bursting is thus important not only from the perspective of grain viability and plant fertilization but also in the context of human health, as ruptured airborne grains can release respirable fragments which can provoke allergic reactions in exposed sensitive individuals [13–15]. In regions of intense agricultural activity, for example, some monocultures—particularly those pollinated by wind—can produce huge quantities of pollen which is prone to bursting [16], and similar problems occur in urban horticultural planning [13, 17]. While grain bursting and consequential release of the pollen grain interior is not characteristic only of allergenic pollen—being a general feature of pollen which hydrates sufficiently rapidly [9]—it is a necessary process for the realization of the allergenic potential of a species [11, 16]. Pollen allergens released in the atmosphere when the grains burst can impact human health for a long time, even when the pollen grains are destroyed and no longer viable [15]. The grains and in particular the cytoplasmic fragments they release when they burst can also act as effective nucleation sites for cloud condensation [18–20] which can influence precipitation [21]. The bursting of pollen grains in the atmosphere is thus relevant both medically and meteorologically. Lastly, the spatial distribution of stresses effected by the inactive mechanical response of the grain could tag the site where the pollen tube originates from and trigger and guide the active mechanisms for its growth [5, 22]. These examples highlight the importance of a quantitative description of pollen grain swelling and bursting in order to understand a wide range of phenomena in different environments and on very different scales.

Any model constructed to represent the influence of changing water content on the pollen grain volume and shape needs to account for the inhomogeneities in the grain wall. Pollen grain walls of an overwhelming majority of gymnosperm and angiosperm species possess discernible regions called apertures, which are known to have chemical and structural composition different from the rest of the wall [3]. The larger part of the pollen shell contains a rigid layer which is mostly made of sporopollenin and is called exine. The apertures are regions in the shell where the exine layer is thinned or entirely absent and mostly consist of cellulose and pectin, making them typically (much) softer than the exine [23, 24]. Apertures thus act as flexible regions in an otherwise stiff pollen shell. As the exchange of water between the pollen grain and its surroundings takes place mostly through the apertures [1, 25], their closure and concomitant infolding of the pollen grain provides a mechanical response which prevents further desiccation and destruction of the grain. It has been recently shown that the success of this process requires a tuned mechanical flexibility of the grain wall, so that the closing of the apertures also pulls in the hard, exine parts of the grain [8, 10] without inducing rupture in the wall material.

Purely mechanical models thus successfully explain the details of infolding and shape changes during the desiccation of pollen grains [8, 10]. Much less is known about the influence of different features of pollen morphology on the swelling and bursting of the grains, which can happen by the rupture of either the apertures or the exine [9]. Our aim is to investigate how pollen grains passively respond to (re)hydration in humid conditions and what role apertures play in this process. In particular, we focus on pollen grains with circular apertures termed *pores* [3], which is the dominant type of pollen of anemophilous (wind-pollinated) plants [26]. Experiments have shown that the pores mechanically respond to hydration [3, 9, 25], yet a theoretical description of this process it still lacking. To describe the swelling and possible rupture of porate pollen grains, we use an elastic model of the grain established previously to describe the shape changes in drying grains [10]. We examine how the presence of the pores influences the swelling of pollen grains as their volume increases due to the influx of fluid from the environment. The increase in grain volume is shown to eventually lead to a rapid inflation of the pores, which results in huge elastic strains in the pore material and most likely leads to the bursting of the pores. We explore how the size, number, and the distribution of pores influence the deformation of the pollen grain and its resistance to bursting. Our results provide quantitative theoretical insight into the inactive mechanisms behind the bursting of pollen grains, and the model we use can be generalized to study the swelling of other types of aperturate pollen grains as well.

## II. RESULTS

### Mechanical properties of porate pollen grains

Pollen species with porate grains are ubiquitous, particularly in anemophilous plants [26], and represent a large proportion of allergenic pollen species [15]. Their pollen grains can have different shapes with a more or less pronounced asphericity. Here, we shall assume that the grains are perfectly spherical in the equilibrium state (thus neglecting any initial asphericity) with an equilibrium radius *R*_0_. Although the model we use allows for different equilibrium grain shapes, the assumption of perfectly spherical grains reduces the number of parameters and allows for a simpler identification and classification of the important features of grain swelling. The porate pollen grains can be further characterized by the number *N* of identical (circular) pores they contain, their distribution, and the angular span of each pore *θ*_0_ (see the examples in Fig. 1).

**FIG. 1.**
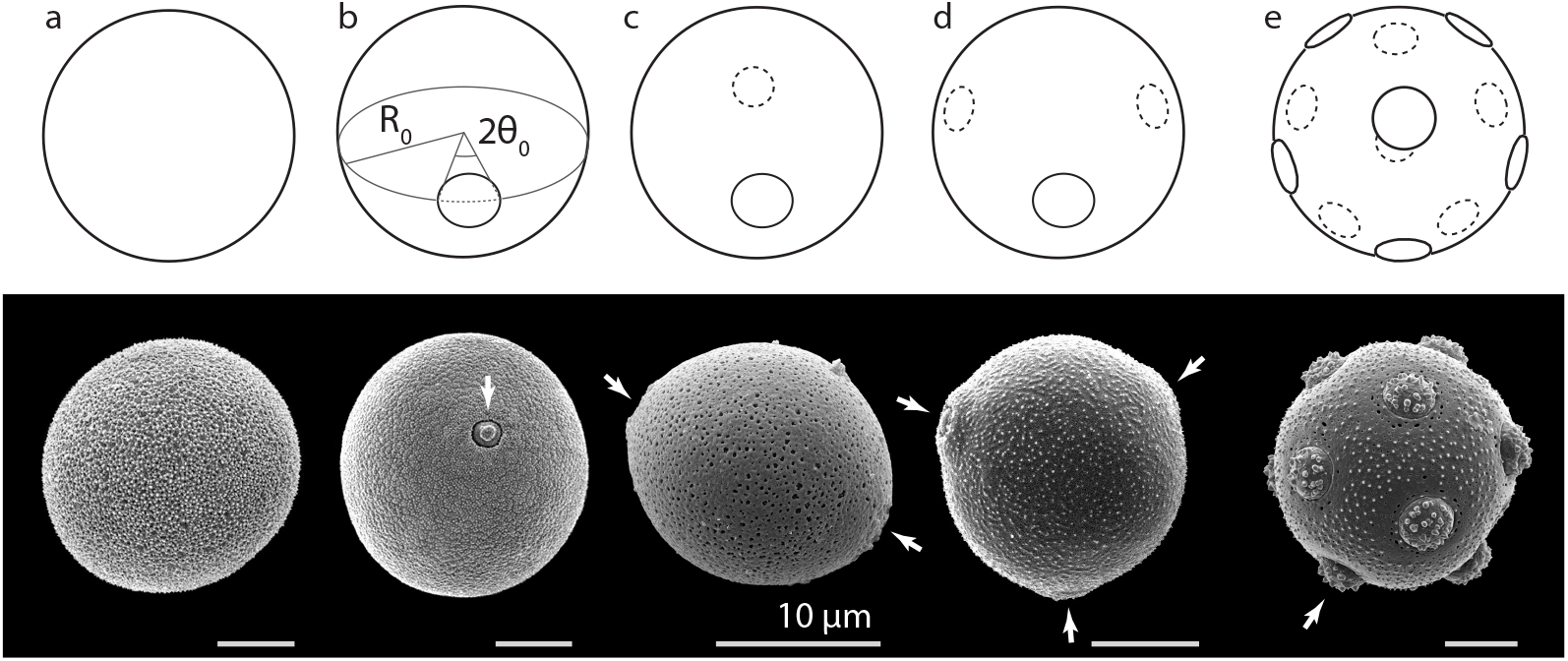
Classes of porate pollen considered in this study: **(a)** inaperturate (*N* = 0), **(b)** monoporate (*N* = 1), **(c)** diporate (*N* = 2), **(d)** triporate (*N* = 3), and **(e)** pantoporate (*N* ~ 10; in this case, *N* = 12). Panel (b) also illustrates the parameters of the pore geometry—its equilibrium, unstrained radius *R*_0_ and the pore opening angle *θ*_0_. Pore size in each schematic porate pollen is *θ*_0_ = 0.2. The bottom part of each panel shows a SEM image of a pollen grain representative of the class: (a) *Populus alba*, (b) *Phleum pratense*, (c) *Besleria solanoides*, (d) *Betula pendula*, and (e) *Stellaria aquatica*. Scale bars in each image represent a length of 10 *μ*m, and the arrows point to the grain pores. In panel (e), only one of the twelve pores is indicated. Pollen images are reprinted with permission from the Society for the Promotion of Palynological Research in Austria; images courtesy of PalDat (2000 onwards, www.paldat.org).

Inaperturate pollen—pollen without any apertures—with *N* = 0 can be considered an extension of the geometric class of porate pollen and represents, at least on the level of our modelling, mechanically the simplest case of a (porate) pollen grain. The genus *Populus*, for instance, contains many anemophilous species with inaperturate pollen, some of which are also moderately allergenic (e.g., *Populus alba*) [27]. Monoporate pollen (*N* = 1) is characteristic of most of the species in *Poaceae* family (grasses) [3], to which some of the most allergenic anemophilous plant species belong (e.g., *Phleum pratense*) [15]. Diporate pollen grains (*N* = 2) with the two pores situated diametrically on the equator of the grain can be found, for instance, in *Morus alba*, although the grains of this species can also have three and four pores. Triporate pollen (*N* = 3) with the three pores arranged equidistantly on the equator of the grain are typical for *Ambrosia artemisiifolia* and the majority of species in the *Betulaceae* family, although some *Betulaceae* species also have five (*N* = 5) equatorially situated pores (e.g., *Alnus glutinosa*). Lastly, *Amaranthus* species are typically pantoporate, i.e., they have many pores (*N* ≈ 20 to 60), distributed nearly uniformly on the grain surface [28]. The opening angles of the pores can be estimated from microscopic images of fully hydrated grains [3] and in general depend on the species, but they are typically about *θ*_0_ ~ 0.1 [29, 30], such as in *Betula pendula* (Fig. 1c) and *Ulmus parvifolia* [3]. In two more extreme examples, the pore angles of *Stellaria aquatica* (Fig. 1e) and *Zea mays* are about *θ*_0_ ≈ 0.18 [31] and *θ*_0_ ≈ 0.05 to 0.07 [32], respectively.

The elastic properties of the pollen grain wall can be roughly parametrized by the Föppl-von Kármán (FvK) number 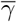 which signifies the relative importance of the stretching and bending energies. The FvK number of pollen grains is typically in the range 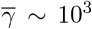 to 10^4^. The pores represent softer regions in the wall compared to the exine part, and we characterize them by a softness parameter *f* (as done previously by Katifori *et al.* [8] and Božič and Šiber [10]) which represents the ratio of the two-dimensional elastic moduli of the pore and the exine, respectively. The ratios of the two-dimensional Young’s moduli and the bending rigidities are thus assumed to be the same, which is not a necessary assumption but conveniently simplifies the parametrization of the problem. The softness parameter of the pores has been estimated in our previous study [10] to be in the range of *f* ~ 0.01 to 0.1. It is a relatively simple feat to measure the volume of a pollen grain as it hydrates by measuring its dimensions [33, 34], and we thus perform elastic calculations at a given volume of the grain, treating it as a mechanical constraint. The shape of the grain is found by minimizing its elastic energy, which also yields the distribution of elastic strains in both the pore and the exine. Details of the elastic model are given in Section IV.

### Swelling of inaperturate pollen grains

As a pollen grain hydrates, its volume increases and its shape changes depending on its elasticity and the distribution, size, and shape of the pores. It is instructive to first consider the swelling of an inaperturate pollen grain—a perfectly homogeneous and spherical elastic shell of radius *R*_0_ enclosing the volume *V*_0_. Such a shell responds to an increase in its interior volume *V* by a simple isotropic increase of its radius *R*. The influence of the presence of pores on pollen grain deformation can then be compared with this idealized case, which approximates a perfectly spherical inaperturate pollen grain (*N* = 0; Fig. 1a).

The extensional strains *ϵ*_0_ in a homogeneous spherical shell are the same everywhere and can be obtained from

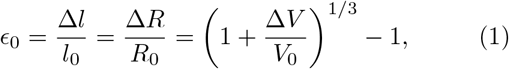

where Δ*l/l*_0_ represents the relative increase of the arbitrarily oriented lengths in the surface of the shell as they extend from *l*_0_ in the initial state to *l*_0_ +Δ*l* in the swollen state [35]. Similarly, Δ*R* denotes the corresponding increase of the radius of the shell when its volume increases by Δ*V* = *V − V*_0_ [36]. For small values of additional volume *v* ≡ Δ*V/V*_0_, Eq. (1) reduces to *ϵ*_0_ = *v/*3. This can be directly applied to the case of inaperturate pollen— once the strain in the exine exceeds a critical value, i.e., once the additional volume *v* becomes large enough, the grain wall will break. For example, if the rupture strain of the exine is 10%, the inaperturate grain can increase its volume by 30% before it ruptures. This estimate is in fact of the correct order of magnitude—rupture strain of ~ 20% has been measured in inaperturate pollen of *Cryptomeria japonica* [37], and similar values are found in experiments on exine deformation and rupture [38]. The estimate effectively presumes that the fracture is brittle and that the material deforms elastically all the way until the fracture. This appears to be a good approximation for both exine and cellulose films [38–40].

### Swelling of monoporate pollen grains

Swelling of inaperturate pollen will lead to exine fracture once the strains in the grain become large enough. If there are pores in the pollen grain, they can relieve some of these strains and thus change the nature of the grain fracture. It is thus of interest to investigate how the presence of a single soft pore (*N* = 1; Fig. 1b) influences the swelling of pollen grains. Fig. 2 illustrates and quantifies this process of an increase in the volume of a pollen grain with *f* = 0.02, *θ*_0_ = 0.15, and 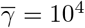. The strain in the pore material is shown in Fig. 2a, which shows the effect of increasing volume on the pore strain averaged over the entire pore surface 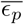 and scaled by the strain which would be characteristic for an inaperturate grain, *ϵ*_0_ of the same volume (i.e., when *f* = 1 or *N* = 0; Eq. (1)). The most notable feature of the monoporate grain hydration is the sudden jump in the pore strain, which for this particular choice of elastic parameters occurs at an additional volume of *v_c_* = 0.263. At this point, the average scaled pore strain increases from 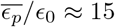 to 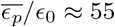. An even more drastic increase is observed in the *maximum* scaled pore strain, which is located at the outermost point of the pore and increases from *ϵ*_*p*;max_*/ϵ*_0_ ≈ 29 to *ϵ*_*p*;max_*/ϵ*_0_ ≈ 196. Note here that this is a *scaled* quantity where *ϵ*_0_ ≈ *v/*3 is about 0.09 at *v_c_* = 0.263, meaning that the maximum pore strain jumps from about *ϵ*_*p*;max_ ≈ 2.6 just before the transition to about *ϵ_p_*_;max_ ≈ 18 after the transition. The sudden transition observed in the (averaged) pore strain exhibits hysteresis, as can be seen by the non-equivalence of the forward and backward minimization paths of gradual increase (hydration) and decrease (desiccation) of internal volume, respectively (marked by arrows in Fig. 2a).

**FIG. 2.**
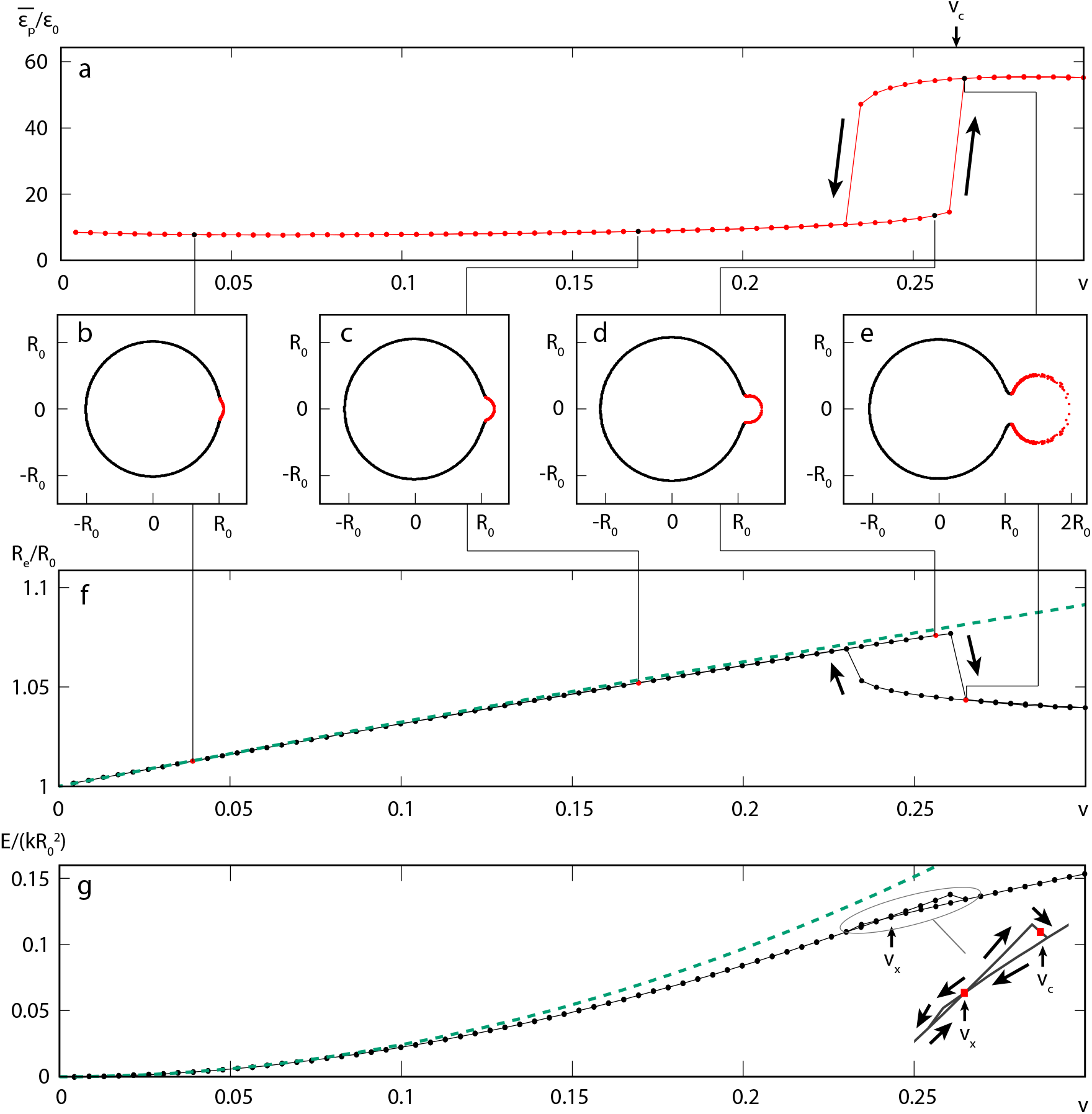
**(a)** Pore strain averaged over the pore area 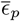 and scaled by strain in a homogeneous grain *ϵ*_0_ (given by Eq. (1)) as a function of additional volume *v* = *V/V*_0_ − 1. **(b)**–**(e)** Cross-sectional projections of the mesh points of the model for *v* = 0.04, 0.17, 0.256, and 0.265. In these panels, the pore and exine materials are denoted by red and black dots, respectively. **(f)** Dimension of the bounding box of the exine part of the pollen grain in the direction perpendicular to the axis connecting the centers of the grain and the pore (i.e., along the ordinate axis in panels (b) to (e)). Dashed line shows the increase in the radius which would occur in an inaperturate, homogeneous grain, *R/R*_0_ = (1 + *v*)^1/3^. **(g)** Total energy of the mesh scaled by 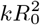 is shown by a black line with symbols. Dashed line shows the continuum limit of the stretching energy in the elastic discrete model of an inaperturate grain, 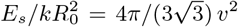. Inset shows the magnified region of *v* where the hysteresis occurs. Mechanical parameters of the pollen grain are *f* = 0.02, *θ*_0_ = 0.15, and 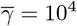 throughout.

Panels (b) to (e) of Fig. 2 show the cross-sectional projections of the triangular mesh of the model grain at different additional volumes, *v* = 0.04, 0.17, 0.256 (just before the transition), and 0.265 (just after the transition). The sudden inflation of the pore (shown in red color) at the critical volume *v_c_* is very pronounced. The mean radius of the exine region of the pollen grain (shown in black color) increases all the way until the transition point at *v_c_* and suddenly decreases afterwards. This change is quantified in Fig. 2f, which shows the effective radius of the pollen grain in the direction perpendicular to the line joining the centers of the grain and the pore (ordinate axis in panels (b) to (e)). Dashed line shows the increase in the exine radius which would be expected in the case of an inaperturate grain, *R_e_/R*_0_ = (1 + *v*)^1/3^. The mean radius of the exine part of the monoporate grain follows this dependence quite closely for small *v*, but the expected increase in *R_e_* slows down as *v* approaches *v_c_* and the exine abruptly *deflates* at the critical volume, compensating in this way for the sudden inflation of the pore.

The sudden inflation of the pore can also be traced in the elastic energy of the pollen grain, shown in Fig. 2g, where a sudden drop in energy is observed at *v_c_*. This calculation also reveals two different energy behaviors which represent two different states of the system. In one of them, the pore still encloses a fairly small volume and has not yet bulged out, whereas in the other, the pore has bulged out and the exine has relaxed. Intriguingly, the two energy behaviors cross at volume *v_x_ < v_c_*, which suggests that the true energy minimum of the system cannot be reached for all volumes, i.e., the pore needs to sufficiently deform before it can bulge out and transit to a lower energy state. This is illustrated in the inset of Fig. 2g, which shows the magnified portion of the forward and backward minimization paths near the intersection of the two energy curves. The inflation of the pore during grain swelling results in lower elastic energies of the grain compared to an inaperturate grain of the same volume. The (stretching) elastic energy of the latter, 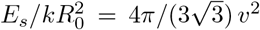, which is the analytical limit of the stretching energy encompassed by the microscopic numerical model in such a situation, is shown by a dashed line.

#### Mechanics of the bursting transition

When the increase in volume of a monoporate pollen grain reaches a critical additional volume *v_c_*, the strains in the pore increase manifold, which can easily cause it to burst. It is therefore appropriate to term the sudden inflation as a *bursting transition*. Bursting of a monoporate grain can be viewed as a transition of the pore through a state when it is maximally curved—this is the state when the inflated pore resembles a hemisphere, Fig. 2d. Up to that point, the pore can resist the internal pressure by increasing its curvature, since the reaction force of the inflated pore is proportional to the inverse radius of curvature (see Appendix A). Past that point, however, the pore radius must increase due to the geometry of the problem, and to resist the additional pressure, the tension in the pore material must increase to resist the same pressure at a larger pore radius. This leads to a sudden inflation of the pore.

The physical mechanism behind the bursting transition can be quantified by equating the normal reaction forces of the exine and the pore at the poles, as they must resist the same internal pressure in the grain [35, 36]. When this analysis is performed for the maximal curvature of the pore, i.e., when the pore attains a (nearly) hemispherical shape (see Appendix A for details of the derivation), one obtains the following equation for the critical additional volume of the grain at which the sudden pore inflation occurs:

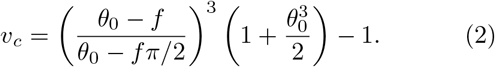

The relation was derived assuming a negligible influence of the bending energies on the transition, which can be seen by the lack of dependence of *v_c_* on 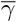 in Eq. (2). When *f/θ*_0_ ≪ 1 and *θ*_0_ ≪ 1, conditions likely to be fulfilled by most pollen grains, the expression for the critical volume reduces to

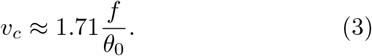

While Eq. (2) and Eq. (3) give only rough estimates of the critical volume of the bursting transition, they are important because they demonstrate that for the mechanical parameters typical for pollen grains, the critical volume should primarily depend on the ratio of the softness of the pore (compared to the exine) and the pore size.

To fully inflate (and eventually burst), the pore must pass through a hemispherical state. Yet even when the pore has not yet sufficiently inflated to reach the hemispherical shape, there might exist inflated states of the pore (i.e. pores larger than a hemisphere) with lower elastic energies. Such states are, however, not geometrically accessible, as they can be reached only after the pore passes through the hemispherical state. In this case, the hemispherical state of the pore represents a geometrical hindrance or a bottleneck and acts as an effective energy barrier. This effect is somewhat similar to the “blow-out” or bursting instability observed in the inflation of flat circular membranes [41, 42]. In our numerical simulations, the energy barrier can be most easily detected by the hysteretic nature of the minimization, as demonstrated in Fig. 2. It is also worth mentioning here that the analogy between pore inflation and the *blowing a balloon confined in a rigid box* has previously been noted by Matamoro-Vidal *et al.* [9] in discussion of experiments on pollen swelling. With this insight, we can now rationalize the two characteristic volumes *v_c_* and *v_x_* which were detected in the dependence of elastic energy of the pollen grain on volume (Fig. 2g). The first of the characteristic volumes, *v_x_*, is the (scaled) additional volume at which the state with the inflated pore (larger than a hemisphere) becomes advantageous with respect to the total elastic energy. The second characteristic volume, *v_c_*, is the critical volume when the internal pressure becomes large enough to surmount the energy barrier of the hemispherical shape and allows the pore to attain the inflated shape. This critical volume, which can be identified in the forward minimization procedure, is relevant for the hydration of pollen grains, during which their volume gradually increases. The analytical considerations given by Eq. (2) also pertain to this volume. The volume where the energy curves cross (*v_x_*) can be detected by combining the results of the forward and backward minimization procedures (see Fig. 2g). At *v* = *v_x_*, there thus exist *two* different states of the grain with the same energy. In one of those states, denoted by 1 in Fig. 3, the pore has not yet reached the critical, hemispherical shape, whereas in the other state, denoted by 2 in Fig. 3, the pore has inflated. These two shapes are obtained in the forward and backward minimization paths at *v* = *v_x_*, respectively.

**FIG. 3.**
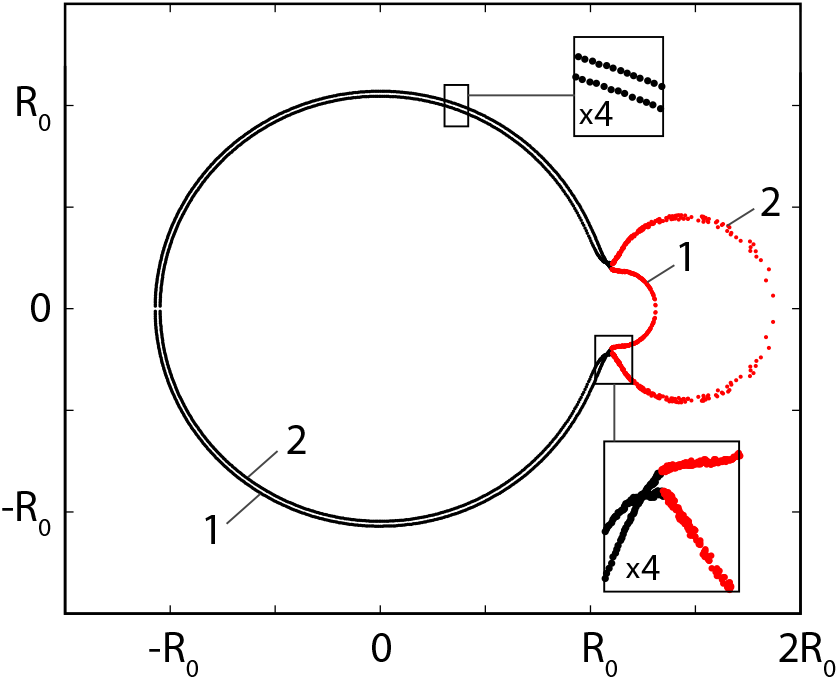
Cross-sectional projections of two states of the grain with the same elastic energy at *v* = *v_x_* = 0.241 (as marked in Fig. 2g). The insets show the magnified regions of the grain cross-sections. Elastic parameters of the grain are *f* = 0.02, *θ*_0_ = 0.15, and 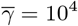.

#### Bursting transition and pore size and softness

To further corroborate the physical interpretation of volumes *v_c_* and *v_x_*, we show in Fig. 4 how the critical volume *v_c_* (squares) and the volume corresponding to the crossing of the energy curves *v_x_* (circles) change with the pore opening angle *θ*_0_ and different values of the pore softness parameter *f*. The analytical prediction of Eq. (2) is shown by full lines, and one can observe that it becomes progressively worse as *f* increases. This is to be expected, since the equation stops making sense as *f* becomes comparable to *θ*_0_—which can happen for either small pore opening angles or sufficiently large *f*—and it has a divergence at *f* = 2*θ*_0_*/π*. Nevertheless, the simple physical reasoning behind Eq. (2) explains the salient features of the numerical results in the range of parameters *f* and *θ*_0_ typical for monoporate pollen grains. It is worth noting that the critical volume at which the pore inflates and bursts does not depend on the elasticity of either the exine or the pore *alone*, but only on the softness of the pore *compared to* the exine (softness parameter *f*). When the pores are sufficiently large (large opening angles *θ*_0_), the energy barrier for pore inflation disappears and the two characteristic volumes become identical, *v_x_* = *v_c_*. The rather rapid inflation of the pore still persists, although it now becomes a continuous phenomenon. Furthermore, the hysteresis of the numerical calculation disappears, which indicates the disappearance of the energy barrier for the bursting transition—for *f* = 0.02, for instance, this happens for opening angles larger than *θ*_0_ ≈ 0.2 (Fig. 4b). The increase in the strains in the pore during the bursting transition becomes smaller as the pore gets larger.

**FIG. 4.**
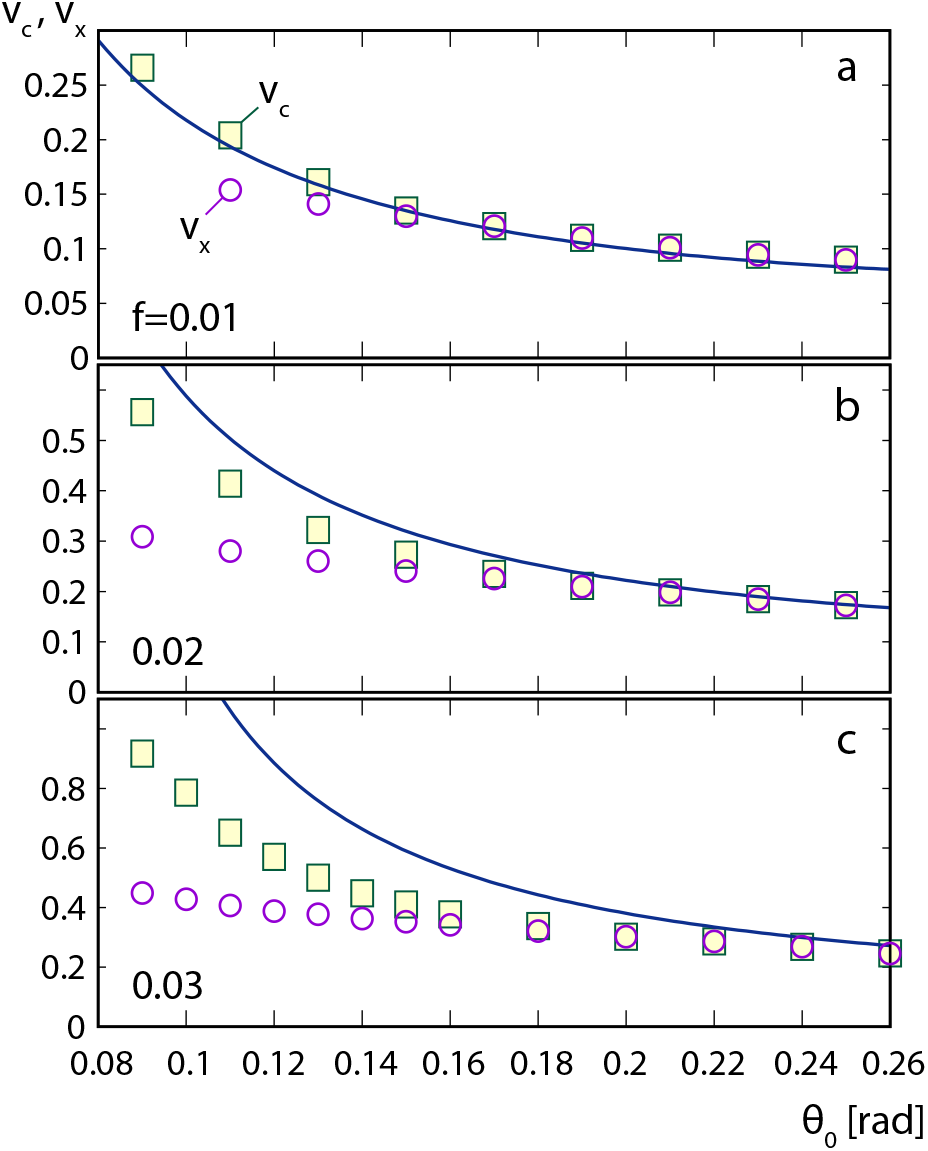
Critical additional volume *v_c_* at which a pore rapidly inflates (squares) and additional volume *v_x_* at which the state with the inflated pore has a lower total energy (circles) as a function of the pore opening angle *θ*_0_ for monoporate grains with 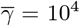. The results are shown for three different pore softness parameters, **(a)** *f* = 0.01, **(b)** 0.02, and **(c)** 0.03. Full lines show the prediction of Eq. (2).

### Bursting of pollen grains with two or more pores

Swelling of monoporate grain eventually leads to an inflation and bursting of the pore, unlike in the case of inaperturate pollen, where swelling causes fracture of the exine once the strains become too large. Pore size and softness set the limit on the amount of swelling a grain can tolerate before it ruptures. Analytical considerations of the bursting transition suggest that the critical volume *v_c_* is not significantly modified by the number of pores in the grain, as long as these are sufficiently small. More precisely, as long as 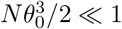, the critical volume of a grain with *N* pores,

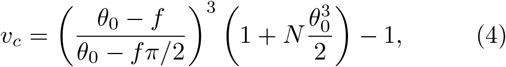

should be essentially the same as in the case of monoporate grain, *N* = 1. One could, however, argue that the final state of the inflated pores is now less strained, as the additional volume is distributed among several pores.

However, numerical calculations indicate that the inflation past the critical point is always asymmetric. Even the slightest difference between the pores leads to a situation in which one of the pores inflates much more than the rest and thus bursts. This is shown in Fig. 5 for a pantoporate pollen grain with *N* = 12 pores, where a single pore suddenly inflates after the bursting transition while the other 11 pores deflate. The surface of the pollen grain is colored in accordance with the local strain in the grain so that the darkest blue color and the brightest yellow color represent the smallest and the largest values of averaged strain when it is larger than it would be in an inaperturate grain with the same additional volume. One can observe that the regions with an increased strain compared to the inaperturate grain are restricted exclusively to the pores. On the other hand, the strains in the exine are quite uniform and reach only about 71% of the value that they would in an inaperturate grain (see the inset of Fig. 5). This demonstrates that the pores act to relieve some of the strain on the exine, the more so the smaller the ratio *f/θ*_0_—the softer and larger the pores are, the more stress on the exine they can relieve (see also Appendix A).

**FIG. 5.**
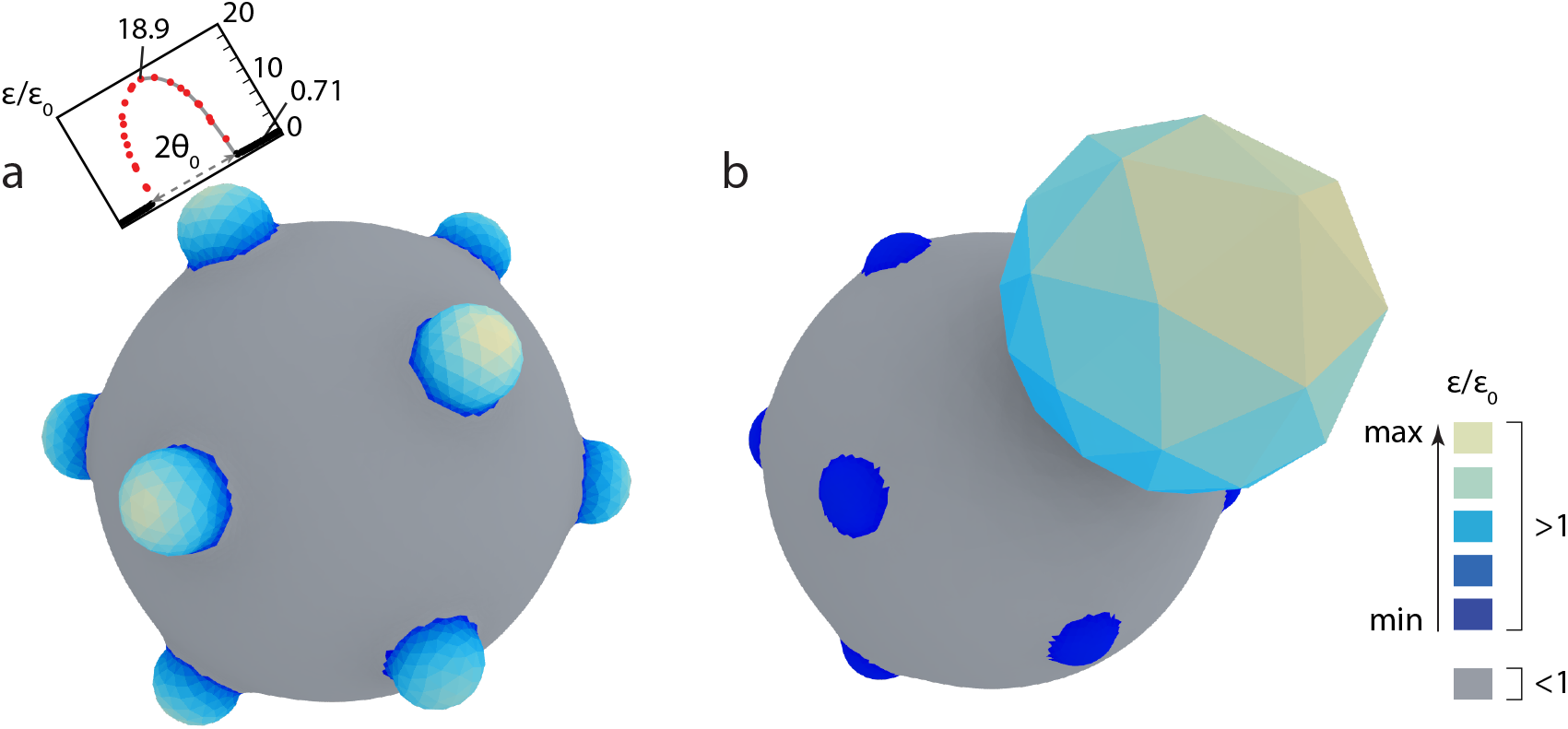
Pantoporate grain with *N* = 12 pores arranged around icosahedron vertices **(a)** right before the bursting transition (*v_c_* = 0.389) and **(b)** just after it (*v_c_* = 0.402). The large sphere-like bulge in the upper right portion of panel (b) is the inflated pore. Mesh triangles are colored in accordance with the scaled local strain *ϵ/ϵ*_0_. The parts of the grain where the strains are smaller than in the inaperturate case (*ϵ/ϵ*_0_ *<* 1) are colored in grey. The parts where *ϵ/ϵ*_0_ *>* 1 are colored using the color-scale shown in the legend so that blue and yellow colors correspond to smallest and largest values of *ϵ/ϵ*_0_ *>* 1, respectively. The maximal relative strains in the grains in panels (a) and (b) are *ϵ/ϵ*_0_ = 18.9 and 178, respectively, in both cases corresponding to the brightest yellow color. Inset in panel (a) shows the cross-sectional profile of the relative strains in the plane which contains the maximally strained point of the pore (its pole) and the grain center. The points corresponding to the pore and the exine are shown by red and black circles, respectively. The x-axis in this diagram is the angular coordinate of the mesh points in the chosen cross-section and is appropriately scaled to match the 3D representation. The elastic parameters of the pollen grain are *f* = 0.02, *θ*_0_ = 0.15, and 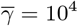.

In Fig. 6, we show the bursting volumes of pollen grains with *f* = 0.02 and *N* = 2, 3, and 4 pores arranged equidistantly along the equator of the grain as well as of a pantoporate grain with *N* = 12 pores arranged on the vertices of an icosahedron. Diporate pollen grains with two diametrically positioned pores behave essentially the same as monoporate grains (Fig. 6a): their critical volumes differ very little, and the two pores in diporate grains do not interact in the relatively wide range of pore opening angles considered. However, as the number of pores increases, critical volumes of such grains begin to deviate from the values obtained in monoporate grains, which can be interpreted as a consequence of an effective elastic interaction between the pores. This effect occurs only for sufficiently large pore opening angles, when the pores themselves become large and approach closer to each other. The region of *θ*_0_ where this effect becomes noticeable is shown by thick light-blue lines on *x*-axes in panels (b) to (d) of Fig. 6. As the number of pores in pollen grains increases, the range of pore sizes where pores interact with each other increases as well, and the pores start to interact at ever smaller values of *θ*_0_. The effect of the pore-pore interaction cannot be accounted for solely by the volume they share, as predicted by Eq. (4), because this provides only a barely visible correction for *N* = 3 and 4 pores (since 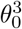 is a rather small quantity). In the case of *N* = 12 pores, Eq. (4) significantly underestimates the correction to the monoporate case, which can be seen by comparing the dashed line in Fig. 6d with the numerically obtained results.

**FIG. 6.**
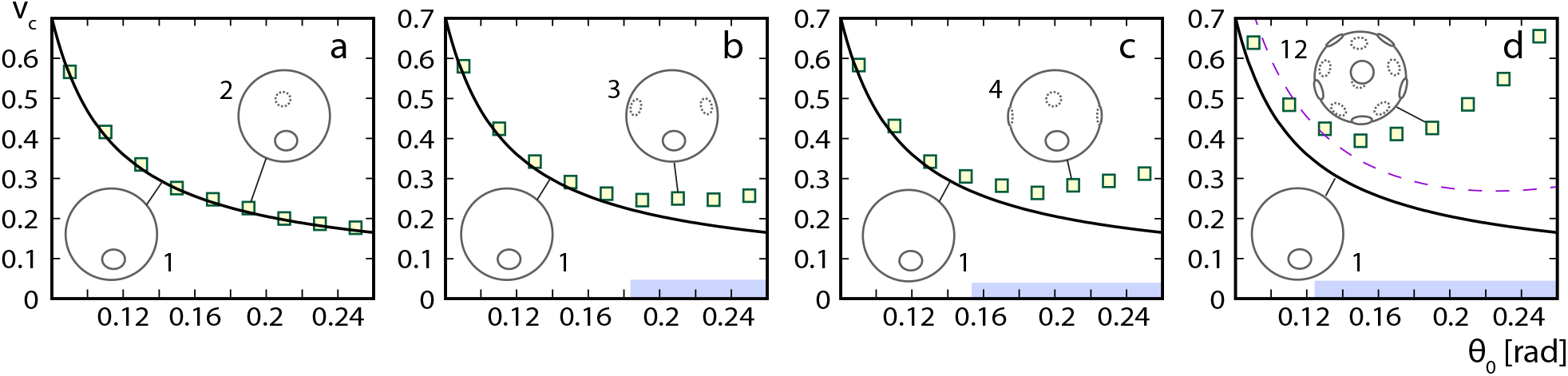
Critical volume *v_c_* for pollen grains with *f* = 0.02 and 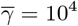 as a function of pore opening angle *θ*_0_ in grains with *N* = 2, **(b)** 3, **(c)** 4, and **(d)** 12 pores. Numerical results are denoted by symbols and full lines represent critical volumes obtained numerically for monoporate grains. Dashed line in panel (d) show the results of Eq. (4) for *N* = 12. Thick light-blue lines above the *x*-axes in panels (b), (c), and (d) indicate approximate regions of *θ*_0_ in which the pores interact with each other.

These discrepancies can be understood by examining the mean distance between the pores. In the case of *N* pores distributed equidistantly along the equator, the pores start to touch each other when *θ*_0_ = *π/N*. At this point, the entire geometry of the problem fundamentally changes and the *N* individual pores merge into a single pore, forming an equatorial poral belt. Such pollen grains with a ring-like aperture at the equator can indeed be found in, e.g., *Zamioculcas zamiifolia* and *Gonatopus angustus* in the *Araceae* family. Geometric aspects of the problem—other than the volume occupied by the pores—are not accounted for by Eq. (4), and the deviation of the critical volume from the monoporate case indicates an elastic interaction between the pores mediated by the exine between them. A physical explanation of this effect requires considerations of the pore packing on the grain surface, their mutual distance and the total area they cover, which is a problem of characteristic lengths and areas rather than volumes. For our purposes, it is important to note that for pore sizes typical for allergenic grains, *θ*_0_ ~ 0.1, the pores can be considered to be independent even when they are numerous, since the critical volumes below about *θ*_0_ ≈ 0.12 are quite similar for *N* = 1 to 4 and even for *N* = 12. In this region, the predictions of Eq. (4) are essentially fulfilled.

## III. DISCUSSION

The mechanical model of the pollen grain adopted in this study predicts that the pores deform significantly more than the exine as the grain swells. The presence of the pores relieves the stress on the exine and reduces it below the values which it would attain if the pores were not present (Fig. 5). At the same time, the pores are also the weak parts of the grain wall which are likely to rupture first once the grain reaches a certain level of hydration, since they undergo a rapid inflation at a critical additional volume of the grain. This pore bursting mechanism, where the pore bulges out and assumes a hemispherical shape just before rupture, has been observed in experiments [13, 43] and is similar to one of the grain rupture mechanisms proposed by Matamoro-Vidal *et al.* [9] for pollen grains with *resistant* exine and *delicate* intine, which in our model corresponds to low values of *f*. Another mechanism observed by Matamoro-Vidal *et al.* [9], a swelling of the grain without either pore or exine bursting and without significant bulging of the pore, is also included in our model which indicates that this might be typical for grains with smaller pores (i.e., small *θ*_0_). The pores are less prone to bursting as they become smaller (see Fig. 4), and since the critical volume of bursting is inversely proportional to *θ*_0_ (Eq. (3)), smaller pores do not undergo the bursting transition until the grain swells to a large extent. In this case this also means that the exine becomes more strained and that it can break before the pores do. Fracture of the exine was also observed by Matamoro-Vidal *et al.* [9], however, only in inaperturate pollen grains with a relatively thin exine. These observations and our model therefore suggest that the pores are, in general, indeed the weak spots of the grain and that they will—if present—typically rupture first.

The critical volumes predicted by our model can be related to values observed in experiments. When the increase in mass of pollen grains was measured at different relative humidities (RH) [20], it was found that until RH ≈ 85%, the grains absorb water internally, while for even larger values of RH, a water layer forms on the external surface of the grains. At RH = 85%, mass of pollen grains increases by about 50%, and this number does not appear to vary much between different pollen types [20]. Maximal volume expansion of pollen grains in the atmosphere can be thus roughly estimated to be about *v* ≈ 0.5, which is similar to the typical values of *v_c_* obtained in our study. This suggest that our model covers the salient aspects of pollen grain swelling and provides a correct estimate of the characteristic energies involved. In particular, the model supports the observation that pollen grains in the atmosphere are in a critical surrounding where changes in humidity may easily lead to grain bursting, depending on the structure of the grains [11]. Grains with sufficiently hard and small pores can sustain a large volume increase without their pores bursting. Although colpate pollen (pollen with elongated apertures) is not the subject of our work, it is nevertheless of interest to note that some colpate pollen, e.g., *Petunia hybrida*, can swell to a huge extent, increasing its volume two or three times upon hydration [33].

Diameters of pollen grains and the size of their pores show a significant correlation across different species of grasses [29, 30], which suggests that their pore opening angles are approximately constant and can be estimated to be in the range of *θ*_0_ ≈ 0.06 to 0.09 (at least in the grass species studied). Such a low value of *θ*_0_ suggests that the pores of pollen grains of grasses are, interestingly, not particularly prone to bursting. This is partially confirmed by experiments [11], where it was found that while 87% of *Betula pendula* pollen grains release the sub-pollen particles when hydrated for 10 minutes, only 40% of *Phleum pratense* pollen grains release the sub-pollen particles under the same conditions. The mean pore opening angle of monoporate grains of *Phleum pratense* (see Fig. 1b) can be estimated to be about *θ*_0_ ≈ 0.08, while the pores of triporate pollen grains of *Betula pendula* have a significantly larger opening angle of *θ*_0_ ≈ 0.13 [3] (see Fig. 1d). This fact alone might thus explain the more frequent rupture of *Betula* pollen in a humid atmosphere.

On the other hand, such simple reasoning does not explain the fact that triporate pollen grains of *Parietaria judaica* with *θ*_0_ ≈ 0.09 [3] burst five times less frequently than pollen grains of *Phleum pratense*, even though they have similar pore opening angles. One should, however, keep in mind that the argument depends on the assumption of a similar value of aperture softness *f* and different values of *f* can be expected in different pollen species [44]—not only because of the differences in apertures and their thickness but also because of the differences in exine thickness and composition, since *f* is a parameter which depends on the *relative* softness of the apertures. While the values of *f* in porate pollen grains are in general unknown, it is nevertheless possible to estimate them in some cases. Rupture of pentoporate pollen grains of *Ulmus parvifolia* has been recorded in a video sequence by Miguel *et al.* [13], and individual frames can be used to determine both *θ*_0_ and *v_c_*, since the volumes right before and after pore rupture can be determined from the sizes of the grain in different frames. This analysis gives *θ*_0_ ≈ 0.11 and *v_c_* ≈ 0.3, which enables one to combine the two numbers and estimate the pore softness to be *f* ≈ 0.015 (see Fig. 4), in line with previous estimates [10].

Pore sizes in pollen of grasses appear to be particularly small, which could signify an evolutionary path which on the one hand allows for a soft spot in the grain to ease the pollen tube growth and on the other hand maximally reduces its size to retain the mechanical consistency of the grain. The minimal pore size is constrained by the by the size of sperm cell which has to pass through the pore into the pollen tube, and a comparison of typical sizes indeed suggests that the pores in grasses are maximally reduced. There, however, also exist pollen grains with a number of quite large pores—such are, for example, the pantoporate grains of *Gypsophila perfoliata* and *Stellaria aquatica* (Fig. 1e). Such an evolutionary solution enables accommodation of large additional volumes before one of the pores bursts. This is manifested by the characteristic shape of the dependence of *v_c_* on *θ*_0_ in Fig. 6d where *v_c_ increases* with *θ*_0_ for sufficiently large pores. This shows that large critical volumes can be obtained not only by a single small pore but also by many large pores (*θ*_0_ *>* 0.16 for the parameters used in Fig. 6). In this respect, it is interesting to note that pantoporate pollen grains appear to have evolved independently many times in different clades of flowering plants [45].

Thinning of the pollen wall in the form of an aperture enables an efficient initiation of the pollen tube growth but at the same time also represents a mechanical weakness of the pollen grain. The same mechanical devices which would in proper conditions aid pollen tube germination can lead to grain rupture and the release of cytoplasm if the grain hydrates in a surrounding with an inadequate osmolarity and ionic content, as might happen in the atmosphere. We have shown that the additional volume *v_c_* which can be sustained by a nearly spherical porate grain and the potential of the grain to rupture are determined predominantly by the ratio *f/θ*_0_ (Eq. (3)), which is a dimensionless parameter combining the pore softness and its size. Other properties of the grain, such as the number of pores, their precise distribution, and the contribution of bending in the process of bursting, appear to be less important, as long as the pores are sufficiently small.

## IV. MATERIALS AND METHODS

### Elastic model of the pollen grain

Construction of the elastic triangular mesh of the pollen grain and the elastic energy functional assigned to it mostly follows the method used previously by Božič and Šiber [10]. We assume that there exists an unstrained state of the pollen grain of volume *V*_0_ which is perfectly spherical and represents the reference, equilibrium state of the problem. The pollen grain can either desiccate, which leads to a reduction of its volume *V < V*_0_, or it can further hydrate, leading to an increase of its volume *V > V*_0_. The second case is of interest to us in this work.

#### Elastic energy of the grain

The elastic energy of the pollen wall can be formulated so that the microscopic energies effectively reside in the edges of the triangular mesh,

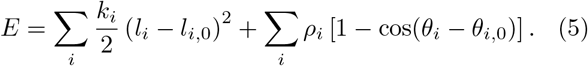

The first and the second term in Eq. (5) are the stretching and the bending energy, respectively. The mesh (triangle) edges *i* have lengths *l_i_*, which, in general, differ from their equilibrium lengths *l_i,_*_0_ in the unstrained state. The stretching energy of an edge is proportional to the square of its extension Δ*l* = *l_i_ − l*_*i*,0_; i.e., each edge *i* acts as a Hookean spring with a spring constant *k_i_*. The bending energy associated with an edge can also be formally tied to the angles *θ_i_* between the two triangle faces which share the edge *i*. The bending energy depends on the difference between the actual and the equilibrium angles *θ_i_* and *θ*_*i*,0_, respectively.

The bending and stretching elastic constants of an edge can take on two different values, depending on whether the edge belongs to the exine or to the pore region of the grain wall, since the two regions have different elastic properties. For edges in the exine region, *k_i_* = *k_ex_* and *ρ_i_* = *ρ_ex_*, while for edges in the pore region, *k_i_* = *k_p_* and *ρ_i_* = *ρ_p_*. The values of the elastic constants in the pore region are scaled by a softness parameter *f <* 1 [8] so that *k_p_* = *fk_ex_* and *ρ_p_* = *fρ_ex_*. Edges which have bounding vertices in different regions, one in the exine and the other in the pore region, are assigned a stretching constant of *k_i_* = (*k_ex_*+*k_p_*)/2. Similarly, when the geometrical centers of two faces sharing an edge are in different regions, the edge is assigned a bending constant of *ρ_i_* = (*ρ_ex_* + *ρ_p_*)/2.

We also introduce a dimensionless quantity 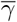 which signifies the relative contributions of the bending and stretching energies and is defined as

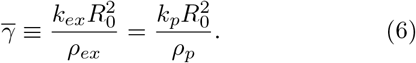

This quantity, used also by Božič and Šiber [10], differs from the Föppl-von Kárman number of the exine (or pore) sheet by a factor on the order of unity [46–48].

#### Mesh triangulation

The equilibrium mesh of triangles is preconditioned to produce the isotropic stains in Eq. (1) as accurately as possible. This is done by optimization of a mesh obtained from marching triangulation [49]. The preconditioning procedure effectively homogenizes the mesh so that the local elastic behaviors are as uniform as possible for a given initial marching triangulation. Once the mesh is optimized, the resulting edge lengths and angles are defined to be the equilibrium values of the edge lengths and angles, and this produces the set of constants *l*_*i*,0_ and *θ*_*i*,0_ in the energy functional in Eq. (5). When such a mesh is inflated for a grain without pores (*k_i_* = *k_ex_* for all edges *i*), the strains are checked to be nearly uniform and isotropic throughout the mesh—as they must be according to Eq. (1). Indeed, they differ from the analytical prediction by at most ~ 1%. For the results shown in this work, the stress- and strain-free state is a spherical shell of radius *R*_0_ = 38.4*a*, where *a* is the mean length of the mesh edge. The mesh has 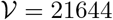 vertices, 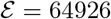 edges, and 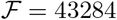 faces, and 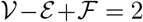, as it must be according to the Euler formula for polyhedra [50].

#### Minimization procedure

The energy functional in Eq. (5) is minimized with respect to the coordinates of the mesh vertices using conjugate gradient method described by Hager and Hongchao [51]. The volume of the mesh is constrained by adding an extra term to the energy functional of the form

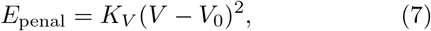

where *K_V_* is the volume penalization constant. The constant must be chosen with care, as too small values do not constrain the volume to a sufficient precision while too large values may result in problems with the conjugate gradient minimization. The appropriate value of the constant *K_V_* can be determined by a sequential minimization procedure, i.e., by a sequence of minimizations in which the constant is multiplied by a factor of *F >* 1 in each step, as described for instance by Šiber [52]. When the minimization is finished, the penalty energy due to the constraint in Eq. (7) must be a negligible percentage of the total elastic energy of the mesh. Once the minimal shape for a given additional volume *v* = *V/V*_0_ − 1 is obtained, all the mesh vertices are randomly jittered, typically by 0.1*a*, and the minimization is repeated. The volume is then increased or decreased, depending on whether the minimization proceeds along the forward or the backward path, respectively.

#### Strains in the pollen grain

To effectively measure the magnitude of the strains in the pollen grain, we define a suitably averaged strain *ϵ_v_* in each vertex of the mesh *v*. The averaged measure of strain is obtained from the area of all the triangles around the vertex *A_v_*,

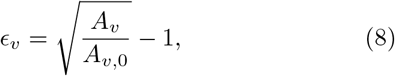

where *A*_*v*,0_ is the area of all the triangles around the vertex *v* in the strain-free state of the mesh. The averaged strain in the entire pore 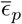 is calculated by summing the strains *ϵ_v_* over all vertices *v* which belong to the pore region and dividing the result by the total number of vertices in the pore.

## ACKNOWLEDGMENTS

AB acknowledges funding from Slovenian Research Agency ARRS (Research Core Funding No. P1–0055).

## Appendix A Analytical approximations for critical volume at bursting transition

Monoporate pollen grain at some additional volume *v* can be approximated as a union of two spherical caps: one, of radius *R_ex_*, representing the exine part of the grain, and the other, of radius *R_p_*, representing the pore. Initially, *R_p_* = *R_e_*, but as *v* increases and the pore inflates, *R_p_* continuously decreases all the way until the pore assumes the shape of a hemisphere, where *R_p_* is the smallest. After this point, any further inflation of the pore requires an increase in *R_p_*.

The internal pressure in the grain *p* is counteracted by the forces in the pollen wall. Examination of the force equilibrium in the two poles of the grain (one in the pore and the other in the exine) when the pore has a hemispherical shape yields

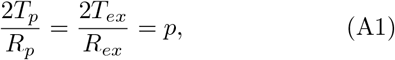

where *T_p_* and *T_ex_* are the tension forces in the pore and the exine, respectively [35]. In the hemispherical state of the pore *R_p_* = *R_ex_θ*_0_ and consequently *T_ex_* = *T_p_/θ*_0_.

If the dominant contribution to the tension forces comes from the stretching part of the elastic energy, the stretching stresses *T* are proportional to strains *ϵ*, which gives

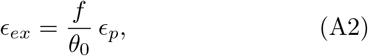

where we have accounted for the fact that the 2D stretching modulus of the pore is softer by a factor of *f* with respect to the corresponding 2D stretching modulus of the exine. The stretching strains can be obtained by examining the distances between the two points in the wall in the stress-free state of the grain and the corresponding distances in the inflated state of the grain, which gives

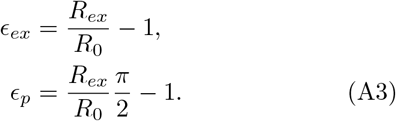

When *θ*_0_ ≪ 1, the scaled additional volume of the pollen grain with a hemispherical pore is given by

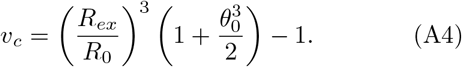

From Eq. (A2) and Eq. (A3), we have

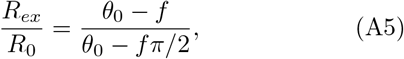

and inserting Eq. (A5) into Eq. (A4), we obtain Eq. (2). For a typical pollen grain, *f/θ*_0_ ≪ 1 and *θ*_0_ ≪ 1, and in this limit we further obtain Eq. (3).

This derivation is only an approximation and is limited in its scope. Eq. (2) predicts that the bursting transition occurs only if *θ*_0_ *> fπ/*2, as *v_c_* → ∞ when *θ*_0_ = *fπ/*2. This is mostly satisfied by pollen grains, but it should be noted that the increase in volume in some pollen grains at bursting is comparable to the volume of a fully hydrated grain, i.e., *v* ~ 1. Furthermore, in the analytical derivation, the characteristic curvatures are assumed to be sphere-like, that is, the same in the two principal directions both in the exine and in the pore. Numerically determined shapes do not have this property, although the principal curvatures do not differ much. The role of the neck region where the pore contacts the exine is also neglected in the approximation, even though it influences the curvatures of the grain shape, even in the polar region of the pore. This is particularly important for larger values of the softness parameter when the approximation of sphere-like curvatures becomes less accurate.

## Validity of analytical approximations for bursting transition

While the numerical results presented in Section II mostly account for the geometry and elastic inhomogeneity which are to be expected in porate pollen grains (parameters *θ*_0_ and *f*), it is of interest to examine the bursting transition for a wider range of parameters. This is because the bursting transition should vanish once the pore becomes hard enough. This will certainly be the case when *f* = 1 and the grain becomes effectively inaperturate, with *v_c_* → ∞. The analytical model in Eq. (2), however, predicts a divergence of *v_c_* when *f* = 2*θ*_0_*/π*. In Fig. 7 we show how *v_c_* changes as a function of *f* for a fixed *θ*_0_ = 0.15. Though large values of *f* (*f >* 0.05) are not expected to be typical for pollen grains [8, 10], the investigation of grains with larger *f* is nevertheless important in the more general context of the mechanics of soft pore inflation. The results indicate that the analytical results serve as a reasonably accurate approximation when *v_c_ <* 0.8, which is a situation typically encountered in pollen swelling and bursting. The approximations become significantly less reliable when *v_c_ >* 1. Somewhat fortuitously, the functional dependence in Eq. (3) can be used as a lower bound estimate for all the volumes *v_c_* studied.

**FIG. 7.**
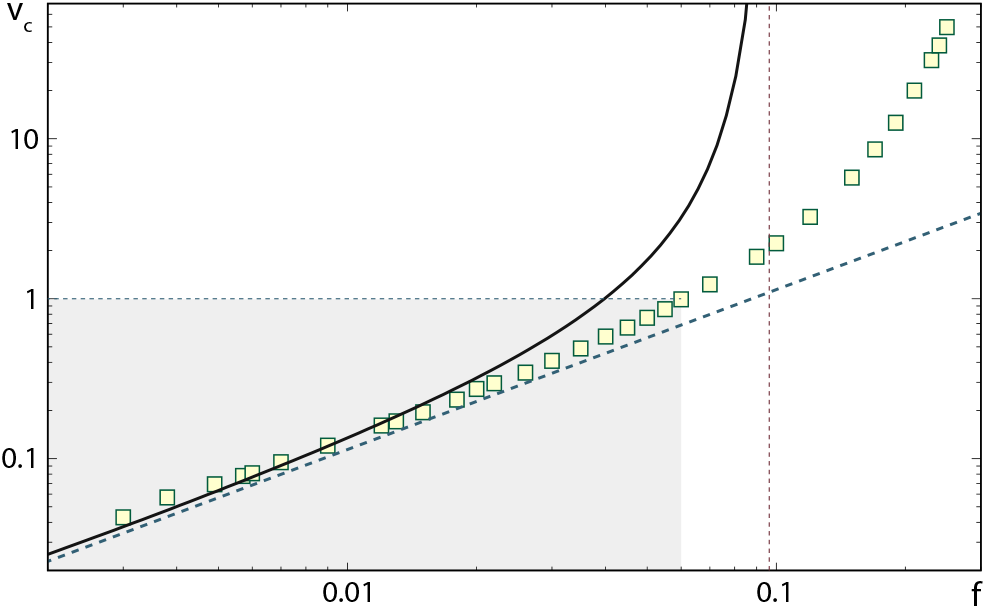
Critical volume at bursting transition *v_c_* for a monoporate pollen grain with *θ*_0_ = 0.15 and 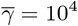 as a function of the pore softness parameter *f*. Numerically obtained results are denoted by symbols. Full and dashed lines are the predictions of Eq. (2) and Eq. (3), respectively. The thin vertical dashed line indicates the value of *f* where the analytical expression in Eq. (2) diverges, *f* = 2*θ*_0_*/π*. The gray area of the plot indicates the range of (*f, v_c_*) parameter space relevant to most pollen grains.

## Internal pressure and bursting transition

The calculations in the main text were performed for a given additional volume of the grain *v*. To each of the grain states obtained with the volume constraint one can also attribute an internal pressure *P*. The internal pressure can be obtained from the normal forces acting on the vertices of the shell in the absence of the volume constraint [36]. Figure 8 shows how the internal pressure in the pollen grain changes as its volume increases—this calculation is performed for the same set of parameters as those used in Fig. 2. A sufficient pressure, denoted by *P_c_* in Fig. 8, is required to overcome the energy barrier for bursting and the pressure drops once the pore inflates. This is similar to the phenomena observed in the inflation of a circular membrane [41] and rubber balloons [53, 54]. Comparing these results with those obtained for thin homogeneous spherical shells by Božič and Šiber [36], we observe that the values of pressure are quite similar at a given increase of volume or exine radius. One can also observe that the presence of a pore decreases the pressure in the grain below the pressure which would act in an inaperturate grain at the same additional volume. This is indicated by the dashed line in Fig. 8, which shows the pressure obtained in the analytical limit of the microscopic model in Eq. (5) for an inaperturat√e grain with the stretching contribution only, 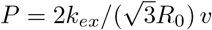 [36].

**FIG. 8.**
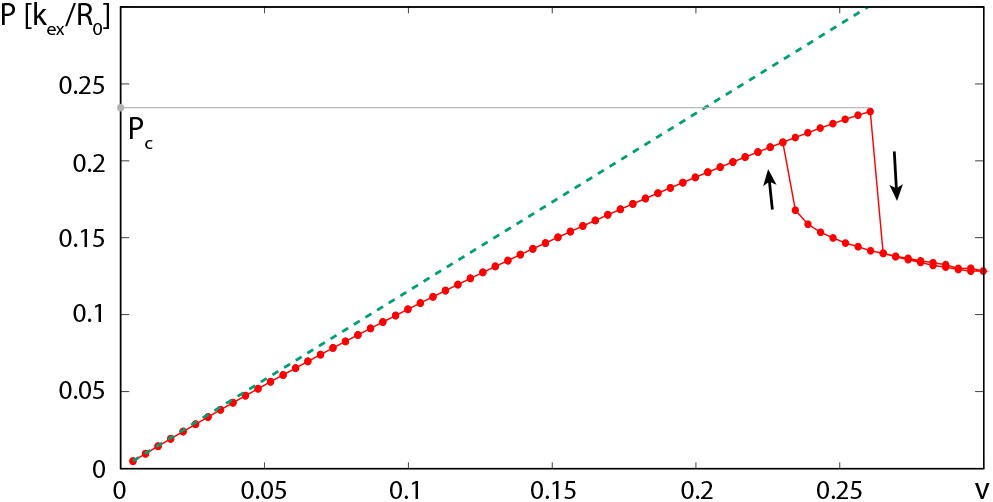
Internal pressure in the grain *P* as a function of the additional grain volume *v*. The pressure is shown in reduced dimensionless units of *kex/R*_0_. Numerically obtained results are denoted by symbols. The dashed line shows the pressure obtained in the analytical limit of the microscopic model for the homogeneous inaperturate shell with stretching energy contribution only, 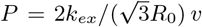. The forward and backward minimization paths are indicated by arrows. The elastic parameters of the calculation are the same as in Fig. 2, *f* = 0.02, *θ*_0_ = 0.15, and 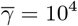.

We have also performed elastic energy minimization at a fixed internal pressure instead of the grain volume. In this case, the energy functional is augmented by an addition of a term *PV* [36, 52], and the volume constraint is released—this approach can be of interest when grain swelling is studied in solutions with different concentrations of non-metabolic sugars [9]. The calculations perfectly reproduce the critical volumes obtained in the fixed volume calculations, but the shapes obtained after the bursting transition (for *P > P_c_*) have very large volumes and are difficult to stabilize in the minimization procedure. The large volumes of these shapes could also have been guessed on the basis of the shape of the pressure dependence in Fig. 8—the pressure *decreases* for *v > v_c_*, which means that the pressure can eventually return to its critical value of *P_c_* only for large values of additional volume *v*. The bursting transition thus figures much more prominently in the calculations at a fixed internal pressure, although the shapes after the transition are difficult to obtain. Both approaches strongly corroborate the necessity of bursting (rupture) of the pore at *P* = *P_c_* or *v* = *v_c_* (*P_c_* = 0.232 *k/R*_0_ and *v_c_* = 0.26 for the case shown in Fig. 8 and Fig. 2).

## Influence of bending energy on bursting transition

All the results shown in the main text are calculated for 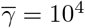. The range of 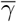 typical for pollen grains was estimated by Božič and Šiber [10] to be between 3000 and 10000. This interval of 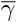 was obtained by requiring that the colpate grains close regularly and completely as they dry up, and might not be entirely relevant for porate grains studied in this work. It is of interest to examine how different values of 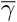 influence the bursting transition. Figure 9 shows how the critical volume at the bursting transition depends on the pore opening angle for three different values of 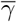. Smaller values of 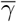 indicate a larger contribution of the bending energy. Figure 9 demonstrates that although the bending energy influences the critical volumes (and pressures), the influence is relatively small and the results obtained in the main text for 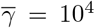 can be considered to be representative for pollen grains. The bursting transition is smoother for smaller 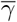, so that the full expansion of the pore occurs in a wider interval of volume.

**FIG. 9.**
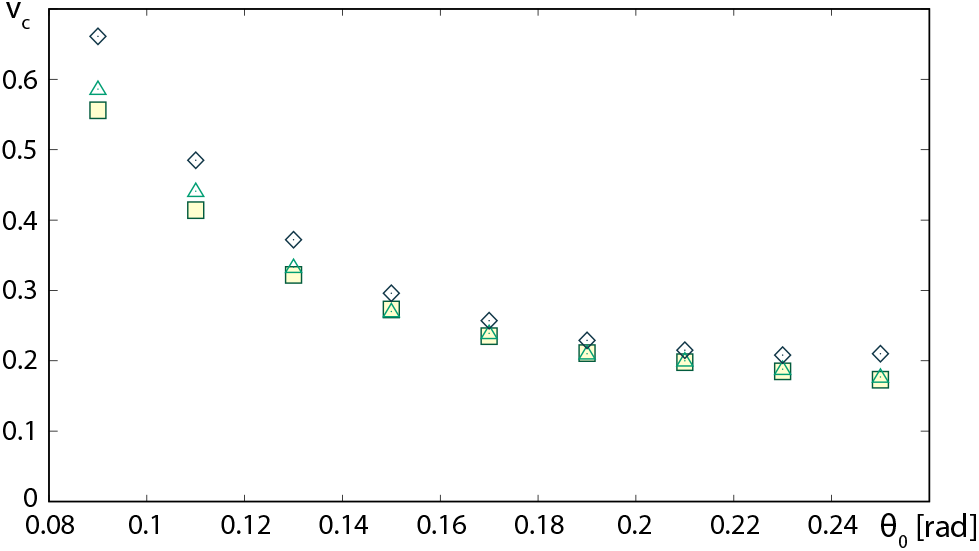
Critical volume at bursting transition *v_c_* as a function of the pore opening angle *θ*_0_ for 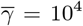 (squares), 5 × 10^3^ (triangles), and 10^3^ (diamonds). The pore softness parameter is *f* = 0.02.

## Nonlinear stress-strain dependence and (non-)universality of bursting transition

The calculated strains in the pore can become huge at the bursting transition. For instance, in the example in Fig. 2, the maximal extension of the pore material *at* the bursting transition is 2.6. Such huge strains may require a modification of the Hookean stress-strain relationship employed in our simulations (see Section IV). While such modifications can in principle be included in our model by allowing the elastic constants to vary with the local strain, in the situation where the elastic responses of the exine and the intine are poorly known such an undertaking does not appear to be particularly enlightening. Nevertheless, it is important to examine the robustness of the predicted bursting transition and investigate whether it persists in different elastic models of the pollen grain. To this end, we modified the stretching part of the Hookean elastic energy (Eq. (5)) to read

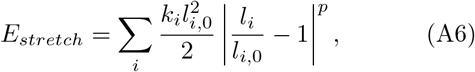

where *p* is a real number, and performed calculations analogous to those shown in Fig. 2 (which correspond to the Hookean elasticity where *p* = 2). The formulation in Eq. (A6) has the advantage that the microscopic stretching and bending elastic constants of the edges *k_i_* and *ρ_i_* are kept the same for all *p* and have the same units as in the Hookean case. The stretching energies of the Hookean case and the *p*-power stretching elasticity coincide in an edge of length *l_i_/l*_*i*,0_ = 2.

The pore bursting transition survives also in the more general parametrization of the stretching energy of Eq. (A6) for a range of values of *p*. There are, however, some differences from the Hookean case with *p* = 2. In particular, while the pores are visibly bulged out at the bursting transition for all powers *p* studied, the shape of the pore at the transition point is inflated a bit over the hemispherical shape when *p >* 2. Furthermore, the critical volumes of the transition sensitively depend on the power *p*, as demonstrated in Fig. 10. For *p >* 2.4, *v_c_* ≳ 1 and the calculations become somewhat irrelevant for most porate grains as they do not tend to swell to such a high degree. It is also possible that for sufficiently large *p*, the bursting transition becomes completely suppressed and never takes place even for infinite increase in volume. This means that the aperture does not importantly modify the strains in the exine and that the entire grain swells almost as if it were inaperturate; this time not because the pore would be too small, but due to the high power *p* in the nonlinear stress-strain relationship. A somewhat similar effect has been noted in relation to the blowout phenomenon of circular membranes and spherical patches, which takes place only for sufficiently low powers *p* in the energy functional of the problem [41].

**FIG. 10.**
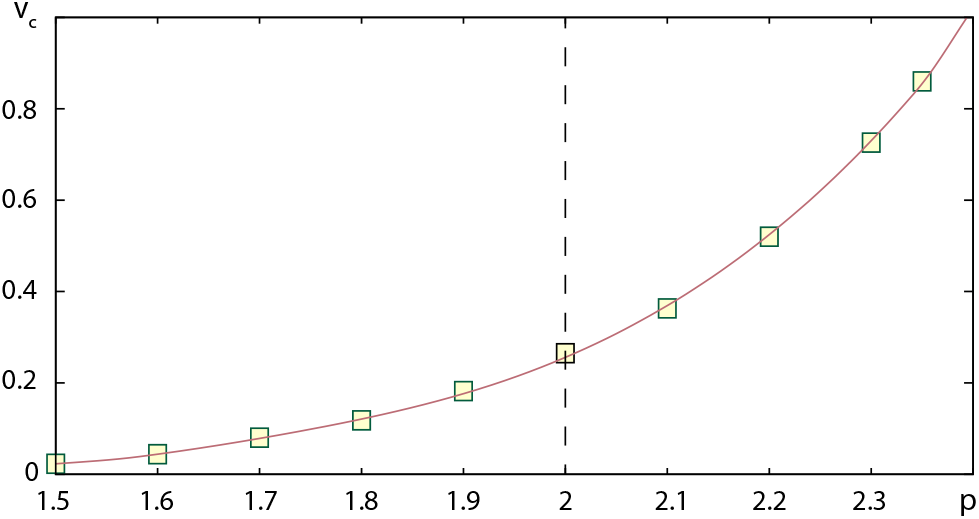
Critical volume *v_c_* at which a pore rapidly inflates as a function of the power *p* in the parametrization of the stretching energy in Eq. (A6). Dashed vertical line emphasizes the point with *p* = 2 (Hookean dependence). The full line is a guide to the eye. The elastic parameters of the monoporate grain are *f* = 0.02, *θ*_0_ = 0.15, and 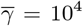

A faithful modelling of pollen grain elasticity would require different parametrizations of the elasticity of the exine and the pores in a much more involved manner than using a simple scaling through a softness constant *f*. This is particularly relevant for large extensions of the exine and the pores, which are likely to be governed by quite different energy functionals and by an effective superposition of several power laws with different values of *p* [55].

